# Dysregulated myogenesis and autophagy in genetically induced pulmonary emphysema

**DOI:** 10.1101/2021.07.08.450201

**Authors:** Joseph Balnis, Lisa A. Drake, Diane V. Singer, Catherine E. Vincent, Tanner C. Korponay, Jeanine D’Armiento, Chun Geun Lee, Jack A. Elias, Harold A. Singer, Ariel Jaitovich

**Author notes:** Correspondence to: Ariel Jaitovich MD, Department of Molecular and Cellular Physiology Albany Medical College, 43 New Scotland Avenue, Albany, NY 12208.

## Abstract

Patients with chronic obstructive pulmonary disease (COPD)-pulmonary emphysema often develop locomotor muscle dysfunction, which is independently associated with disability and higher mortality in that population. Muscle dysfunction entails reduced muscle mass and force-generation capacity, which are influenced by fibers integrity. Myogenesis, which is muscle turnover driven by progenitor cells such as satellite cells, contributes to the maintenance of muscle integrity in the context of organ development and injury-repair cycles. Injurious events crucially occur in COPD patients’ skeletal muscles in the setting of exacerbations and infections which lead to acute decompensations for limited periods of time after which, patients typically fail to recover the baseline status they had before the acute event. Autophagy, which is dysregulated in muscles from COPD patients, is a key regulator of satellite cells activation and myogenesis, yet very little research has so far investigated the mechanistic role of autophagy dysregulation in COPD muscles. Using a genetically inducible murine model of COPD-driven muscle dysfunction and confirmed with a second genetic animal model, we found a significant myogenic dysfunction associated with a reduced proliferative capacity of freshly isolated satellite cells. Transplantation experiments followed by lineage tracing suggest that an intrinsic defect in satellite cells, and not in the COPD environment, plays a dominant role in the observed myogenic dysfunction. RNA sequencing analysis of freshly isolated satellite cells suggests dysregulation of transcripts associated with control of cell cycle and autophagy, which is confirmed by a direct observation of COPD mice satellite cells fluorescent-tracked autophagosome formation. Moreover, spermidine-induced autophagy stimulation leads to improved satellite cells autophagosome turnover, replication rate and myogenesis. Our data suggests that pulmonary emphysema causes a disrupted myogenesis, which could be improved with stimulation of autophagy and satellite cells activation, leading to an attenuated muscle dysfunction in this context.

## Introduction

Patients with chronic obstructive pulmonary disease (COPD) often develop locomotor skeletal muscle dysfunction which entails muscle atrophy and lower force-generation capacity [1, 2]. COPD-associated muscle dysfunction, which occurs more often in emphysema than in chronic bronchitis phenotype [3], is strongly associated with higher mortality and other poor outcomes in these patients [4-6]. Inferential models indicate that these associations persist after multivariably adjusting for the level of pulmonary disease and other covariables suggesting that muscle dysfunction could partially contribute to the worse prognosis [1, 2].

In general, muscle function/force-generation capacity is determined by total mass and fiber’s metabolic properties [1]. Myogenesis, which is muscle turnover driven by progenitor cells such as satellite cells [7], contributes to the maintenance of muscle mass and metabolic profile in the context of organ development, hypertrophy and injury-repair cycles [8-10]. Injurious events crucially occur in COPD patients skeletal muscles in the setting of exacerbations and infections [11, 12], which lead to acute decompensations for limited periods of time after which patients typically fail to recover the baseline status they had before the acute event [13, 14]. Indeed, the frequency and severity of COPD exacerbations and infections powerfully associate with loss of muscle and lung integrity, and with higher mortality over time [13-16]. While strong evidence indicates that dysfunctional myogenesis contributes to muscle loss in non-primarily muscular conditions such as aging and cancer [17-20], recent clinical observations also suggest its role in COPD [18, 21-27]. Moreover, accumulating evidence indicates that biological signals that are prevalent in COPD patients such as hypoxia, hypercapnia and smoking regulate muscle response to injury as well [28-30]. So far, almost no mechanistic research focused on skeletal myogenesis has been conducted on an animal model of COPD-induced skeletal muscle dysfunction. Such information could accelerate research aimed at improving muscle recovery in the setting of acute injurious events following COPD decompensation, with potential mortality benefits.

Autophagy, a major cellular homeostatic mechanism activated by environmental stress [31], contributes to myogenesis by supporting bioenergetic demands for satellite and other stem cells activation [32-35], and by preventing quiescence-to-senescence transition [17, 20, 36]. While consistent clinical observations have revealed a profound autophagy dysregulation in skeletal muscles from COPD patients [37-39], there is no clarity regarding the contribution of abnormal autophagy to COPD muscles integrity. Moreover, while impaired autophagy is known to lead to muscle loss [37], excessive autophagy appears to contribute to muscle wasting as well [40]. So far, no mechanistic research has addressed the role of autophagy in the regulation of satellite cell-mediated myogenesis in the COPD setting.

Mechanistic research focused on COPD-driven locomotor muscle dysfunction requires an animal model that ideally should fulfill the following conditions: 1) be inducible, in order to minimize temporal confounders such as muscle development and age-related sarcopenia [41]; 2) be robust enough, to reminisce the disease severity shown by the majority of COPD patients with muscle dysfunction [42]; 3) develop the muscle phenotype after, and not simultaneously with, the occurrence of pulmonary disease to reflect a secondary COPD comorbidity; 4) recapitulate multidimensional features observed in patients such as morphologic, metabolic and functional aspects of muscle dysfunction [1, 43, 44]; and 5) occur in the context of a pulmonary disease phenotype with airways obstruction, typical histologic changes and other features demonstrated by humans [45, 46].

In the present work, we took advantage of a recently established murine model of pulmonary emphysema-induced skeletal muscle dysfunction to interrogate the animal’s myogenic response. That animal, which highly reminisces the skeletal muscle phenotype demonstrated by COPD patients [43-45, 47, 48], develops pulmonary emphysema in a transgene-driven inducible fashion, allowing control of the time-sensitive myogenic effects of chronic pulmonary disease. This animal model deliberately does not involve cigarette smoking given that this signal has been recently shown to influence muscle response to injury [28], which could confound the specific effect of pulmonary emphysema on myogenesis. Moreover, features characterizing muscle dysfunction do not occur in this animal until the emphysema is established at 8 weeks post induction [43, 44], suggesting a similar trajectory to the demonstrated by COPD patients with muscle dysfunction [1]. Additionally, the timing of induction can be finely controlled to avoid influences of developmental myogenesis and to also develop a robust phenotype in a young adult mouse allowing us to dissociate any age-related effects on reduced myogenic potential from those that are specific to the COPD phenotype. Our central hypothesis was that autophagy dysregulation contributes to inadequate muscle repair in COPD and that autophagy stimulation would attenuate that deficit.

## Results

### Inducible pulmonary emphysema causes muscle dysfunction and an autophagy dysregulation signature reminiscent of COPD patients

To define whether our selected animal model of genetically driven pulmonary emphysema/COPD is an adequate platform to investigate the influence of autophagy on skeletal myogenesis, we provided doxycycline to induce *IL13*^*TG*^ (COPD) mice and compared it with similarly treated *IL13*^*WT*^ (wild type) counterparts. Lung macroscopy demonstrated evidence of hyperinflation as reflected by larger organ size; which was corroborated by microscopic examination of lung sections, which showed evidence of airspace enlargement in *IL13*^*TG*^ compared with *IL13*^*WT*^ animals (**Figure 1-A and B**). Oxygen saturation at room air demonstrated that *IL13*^*TG*^ mice are hypoxemic as opposed to *IL13*^*WT*^ animals which are not (**Figure 1-C**). Moreover, muscle force-generation capacity as measured by the *in-vivo* grip strength test and *ex-vivo* isolated extensor digitorum longus (EDL) muscle contractility test demonstrated evidence of skeletal muscle dysfunction in *IL13*^*TG*^ mice relative to wild type counterparts (**Figure 1-D and E**). Surrogates of muscle atrophy such as muscle mass and individual fibers’ cross-sectional area (CSA) were also significantly reduced in *IL13*^*TG*^ compared with *IL13*^*WT*^ animals (**Figure 1-F and G**). To define whether muscles from COPD mice demonstrate features of autophagy dysregulation, we interrogated a set of genes which altered expressions have been previously reported in analyses of COPD patients muscle biopsies [37]. *IL13*^*TG*^ (COPD) mice demonstrated a similar transcript profile to the previously found in human studies (**Figure 1-H**, see also reference [37]). These data supported the rationale to further investigate autophagy’s effect on myogenesis using validated methods.

**Figure 1.**
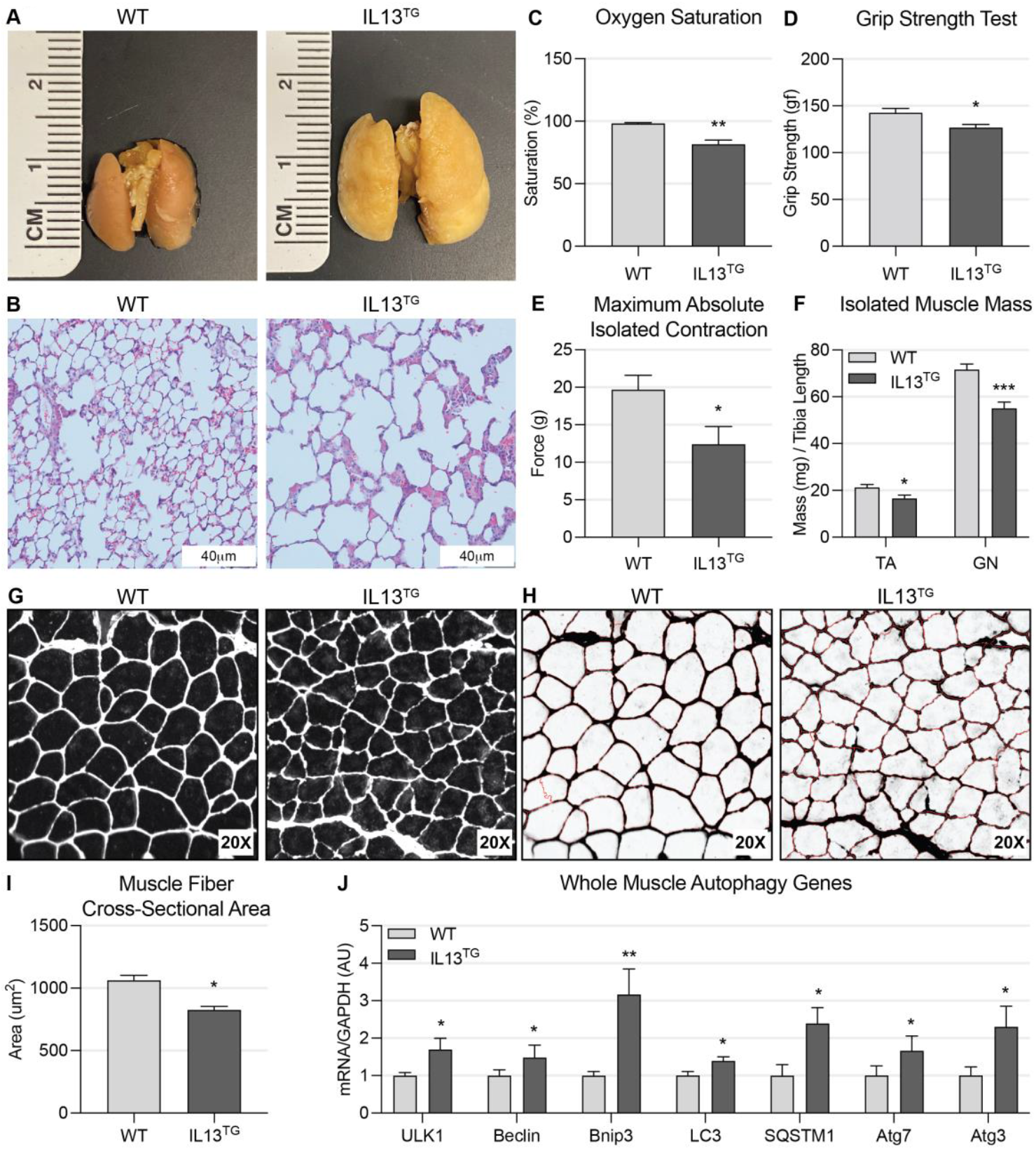
Transgenic *IL13^TG^* mice develop pulmonary emphysema followed by skeletal muscle dysfunction. **A:** Lungs procured from fully induced animals exhibit significant size increase consistent with hyperinflation. **B:** Hematoxylin and eosin staining of lung sections under magnification show damaged alveoli with increased size and irregular shape. **C:** Arterial hemoglobin oxygen saturation at room air is reduced in *IL13^TG^* mice (n=5). **D:** Emphysematous mice demonstrate reduced maximal force-generation capacity as measured by four-limb grip strength (n=5). **E:** Isolated extensor digitorum longus (EDL) muscle stimulated maximum contraction is reduced in *IL13^TG^* mice (n=8). **F:** Freshly isolated wet muscle mass of both tibialis anterior (TA) and gastrocnemius (GN) muscles is reduced in *IL13^TG^* mice (n=8). **G:** EDL muscle sections from *IL13^TG^* animals demonstrate reduced fibers cross-sectional area (CSA) as measured automatically by CellProfilerTM software (n=5). **H:** CellProfiler™ output image files used for unbiased fibers CSA measurement show high fidelity with minimal methodological noise (n=5). **I:** Average fibers CSA is significantly reduced in *IL13^TG^* animals when compared with wild-type (WT) age-matched litter mates (n=5). **J:** Real-time PCR panel of autophagy genes in TA muscle shows a dysregulation profile similar to previous clinical data (n=8); \**P <* 0.05; \*\**P <* 0.01; \*\*\**P <* 0.001.

### Pulmonary emphysema causes suboptimal myogenic capacity

To investigate the myogenic response of *IL13*^*TG*^ (COPD) mice in comparison with *IL13*^*WT*^ (wild type) counterparts, we conducted a two-wave barium chloride (BaCl_2_) injury-repair assay [32]. This two wave injury-repair assay has been previously found to improve the resolution of satellite cell contribution to myogenesis [49]. At sequential timepoints after the second injury, animals were sacrificed and tibialis anterior (TA) muscle histology was analyzed. COPD mice demonstrated a more disorganized early repair compared with wild type littermates. Two weeks post-injury both COPD and wild type mice demonstrated recovery of fibers integrity however, the cross-sectional area of the COPD mice was significantly reduced in comparison with the wild type littermates (**Figure 2-A**). Satellite cells contribute majorly to muscle repair after injury [7]. In order to participate in myogenesis, satellite cells undergo a few cycles of “symmetrical” cell division that maintain their undifferentiated phenotype, followed by an “asymmetrical” division that commits them to the myogenic lineage [8]. To score the replicative capacity of satellite cells isolated from COPD versus wild type mice, we conducted a 5-ethynyl-20-deoxyuridine (EdU) incorporation assay [32]. This method, which is based on EdU incorporation to the DNA every time cells undergo a cycle, has been previously calibrated to resolve satellite cells replicative capacity at 40 hours after their isolation [50]. We observed that satellite cells isolated from COPD mice demonstrated a significantly lower replication capacity in comparison with wild type mice (**Figure 2-B**). To rule out the possibility that the overexpression of IL13 in our COPD mouse model contributed to the dysfunctional satellite cell replication independently of COPD *per se*, we used a second established animal model of COPD based on interstitial collagenase (MMP1) expression [51]. These *MMP1*^*TG*^ (COPD) mice, which also develop features of muscle dysfunction (**Supplementary figure S1**), demonstrate a reduced EdU incorporation in comparison with *MMP1*^*WT*^ (wild type) counterparts (**Figure 2-C**). As these *MMP1*^*TG*^ mice data validated the concept of dysfunctional satellite cell replication in COPD, the following experiments were conducted with *IL13*^*TG*^ and *IL13*^*WT*^ mice, which served as the primary model in this research.

**Figure 2.**
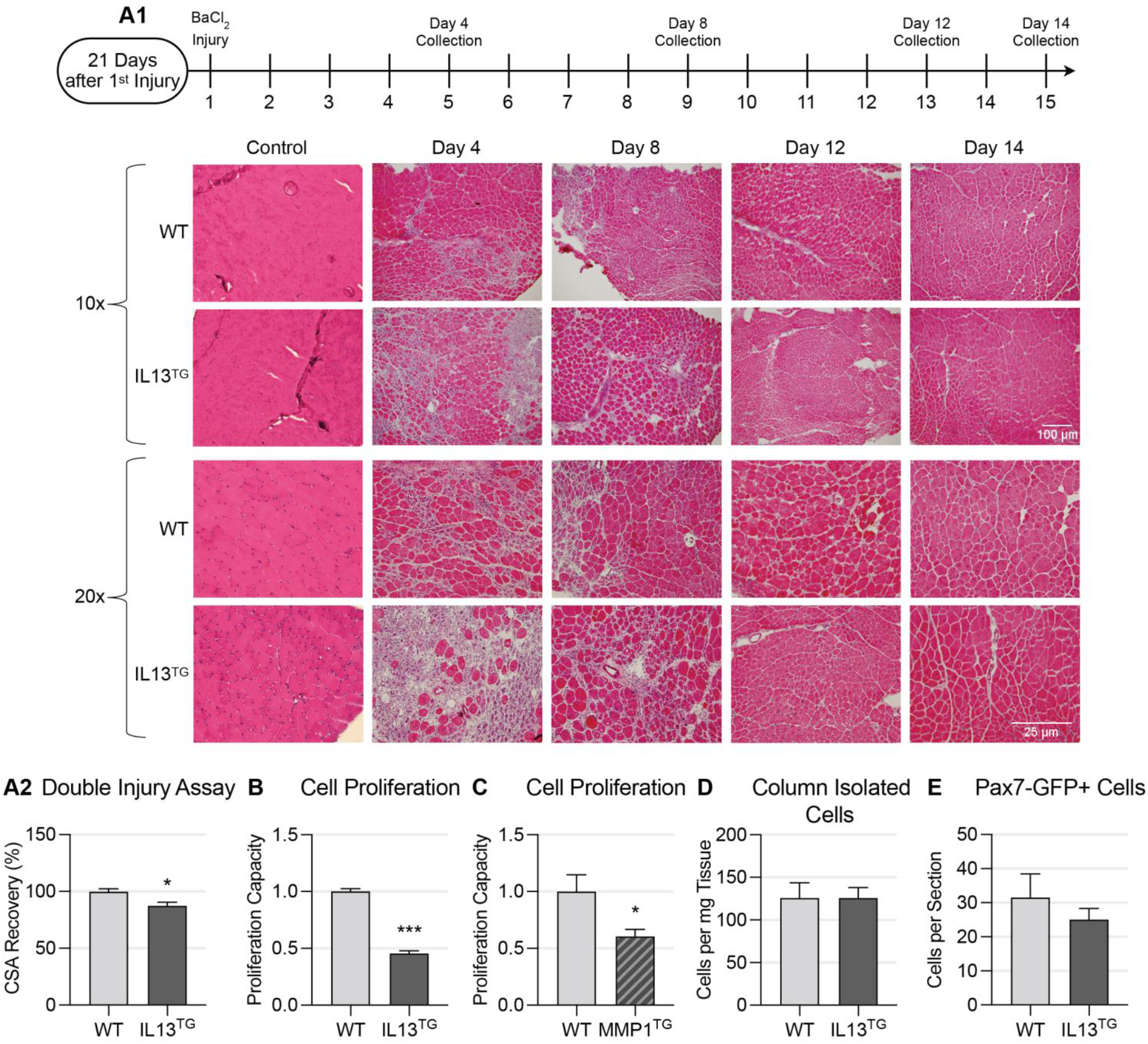
Emphysematous (COPD) mice demonstrate dysfunctional myogenesis. **A1:** Double-injury repair assay reveals suboptimal and delayed muscle repair of *IL13^TG^* over time when compared with WT littermates. **A2:** *IL13^TG^* TA muscle recovery at day 14 as determined by fibers CSA remains significantly below baseline muscle CSA when compared to the individuals counter-lateral leg TA muscle CSA (n=3). **B:** Fully induced *IL13^TG^* animals have significantly reduced myogenic cell proliferation capacity as evaluated by 5-Ethynyl-2′-deoxyuridine (EdU) assay (n=4). **C:** Emphysematous *MMP1TG* mice show a significant reduction in myogenic cell proliferation (n=4). **D:** Integrin α 7 positive cells isolated using MACS cell separation technique showed similar counts between *IL13^TG^* and *WT* animals (n=4). **E:** Pax7-GFP auto-fluorescent muscle stems cells counted in situ using automatic cell counting showed no significant difference between IL13TG and WT mice (n=4); \**P <* 0.05; \*\*\**P <* 0.001.

To define whether the COPD phenotype is associated with a reduced number of satellite cells, we quantified epitope-specific (α7-integrin positive)-isolated cells [52], which were not significantly different between COPD and wild type phenotypes (**Figure 2-D**). Given that isolation of satellite cells disrupts their niche and causes activation [8], we also scored the number of quiescent satellite cells in COPD and wild type animals. As the transcription factor Pax7 is a canonical marker of quiescent satellite cells and is required for their normal function [49], we conducted an unbiased analysis of fluorescent Pax7 positive cells in uninjured muscle sections. To do that, we crossed the COPD and wild type mice with an animal holding an inducible green-fluorescence protein (GFP) reporter in satellite cells (IRES-CreERT2 cassette) inserted after the endogenous termination codon of Pax-7 gene [53]. This model, which captures about 95% of Pax7 positive cells [53], was unbiasedly analyzed accounting for more than 1500 fibers per section. Fluorescent Pax7-positive cells numbers were not significantly different between COPD and wild type mice (**Figure 2-E and supplemental figure S2**). Taken together, these data suggest a suboptimal myogenic response in COPD animals, which is associated with a replicative dysfunction, but not reduced number, of satellite cells.

### COPD leads to an intrinsic dysfunction of satellite cells

To define whether the described myogenic dysfunction is driven by an intrinsic satellite cells deficiency or instead by environmental factors determined by the COPD milieu, we conducted transplantation experiments with lineage tracing assays. To do that, we generated COPD and wild type mice constitutively expressing red fluorescence protein (RFP) in skeletal muscle and satellite cells. These animals, resulting from the crossing of *IL13*^*TG*^ and *IL13*^*WT*^ mice with an RFP-expressing mouse under the control of the β-actin promoter [54], were used to obtain similar numbers of satellite cells, which were then freshly transplanted into receiving animals with either COPD or wild type phenotype (**Figure 3-A**). Given that freshly isolated satellite cells contribute to muscle repair by fusing with preexisting mature fibers or with other satellite cells [8], the RFP expressing fibers after muscle repair in this model are a specific surrogate of transplanted satellite cells-driven myogenesis [55]. Two weeks after transplantation, mice were sacrificed and RFP-positive fibers were counted, indicating that *IL13*^*WT*^-derived cells were able to contribute to larger number of fibers even when transplanted into a COPD animal, whereas *IL13*^*TG*^-derived satellite cells participated in neofibers formation in a substantially lower number even if transplanted into a healthy animal (**Figure 3-B**). These data suggest that cells donated from COPD animals carry an intrinsic defect that could contribute to dysfunctional myogenesis independently of the COPD muscle environment.

**Figure 3.**
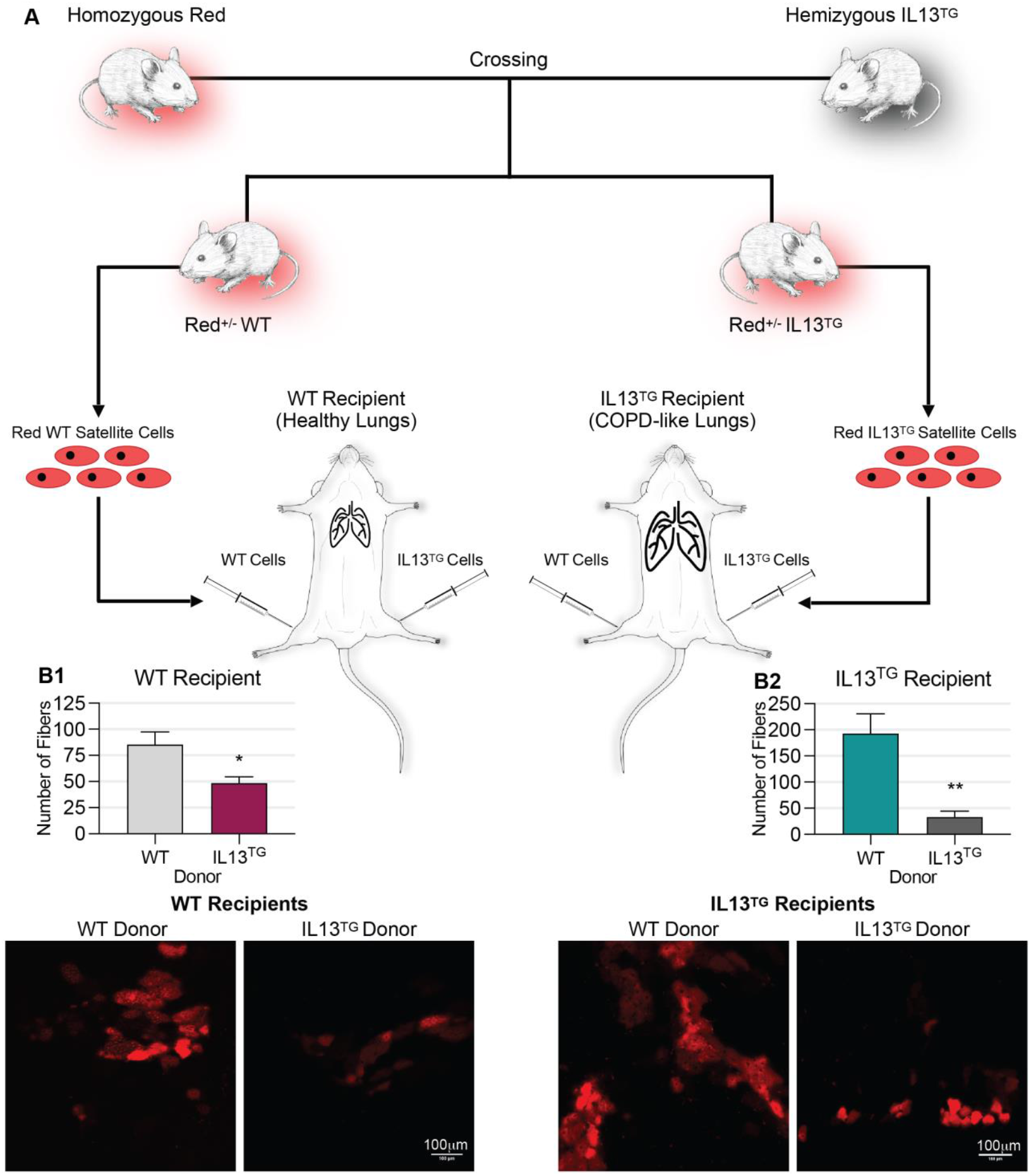
Transplantation experiments with linage tracing reveal that cells donated from COPD mouse donors have altered myogenic capacity. **A:** Cartoon illustrates transplant experimental design. **B1:** *WT* recipients receiving donation of *red-IL13^TG^* cells in one TA muscle and *red-WT cells* in the counter-lateral TA muscle show significantly reduced number of red fibers after repair in *IL13^TG^* receiving leg when compared to the animals counter-lateral *WT* receiving leg (n=4). **B2:** Similar experiments using *IL13^TG^* recipients reveal similar significant reduction (n=5); \**P <* 0.05; \*\**P <* 0.01.

### Transcriptomic profile of COPD mice satellite cells suggests dysregulation of multiple genes associated with cell division

To further interrogate the possible mechanisms of satellite cells dysfunction in COPD, we freshly isolated post-injury cells and sorted them with fluorescence-activated cell sorting (FACS), as previously established [56]; and processed them for RNA sequencing analysis. This analysis revealed that, out of the transcripts captured by the analysis, about 75% of the genes belonging to cell cycle (GO:0007049) and cell division (GO:0051301) ontology terms were dysregulated in COPD in comparison with wild type littermates. Moreover, given that autophagy plays a central role in satellite cells activation and symmetrical cell division [32, 33], we specifically queried for this process and found that about 85% of the genes belonging to autophagy (GO:006914) and regulation of autophagy (GO:0010506) were downregulated in our analysis (**Figure 4 and supplementary table 1-2**).

**Figure 4.**
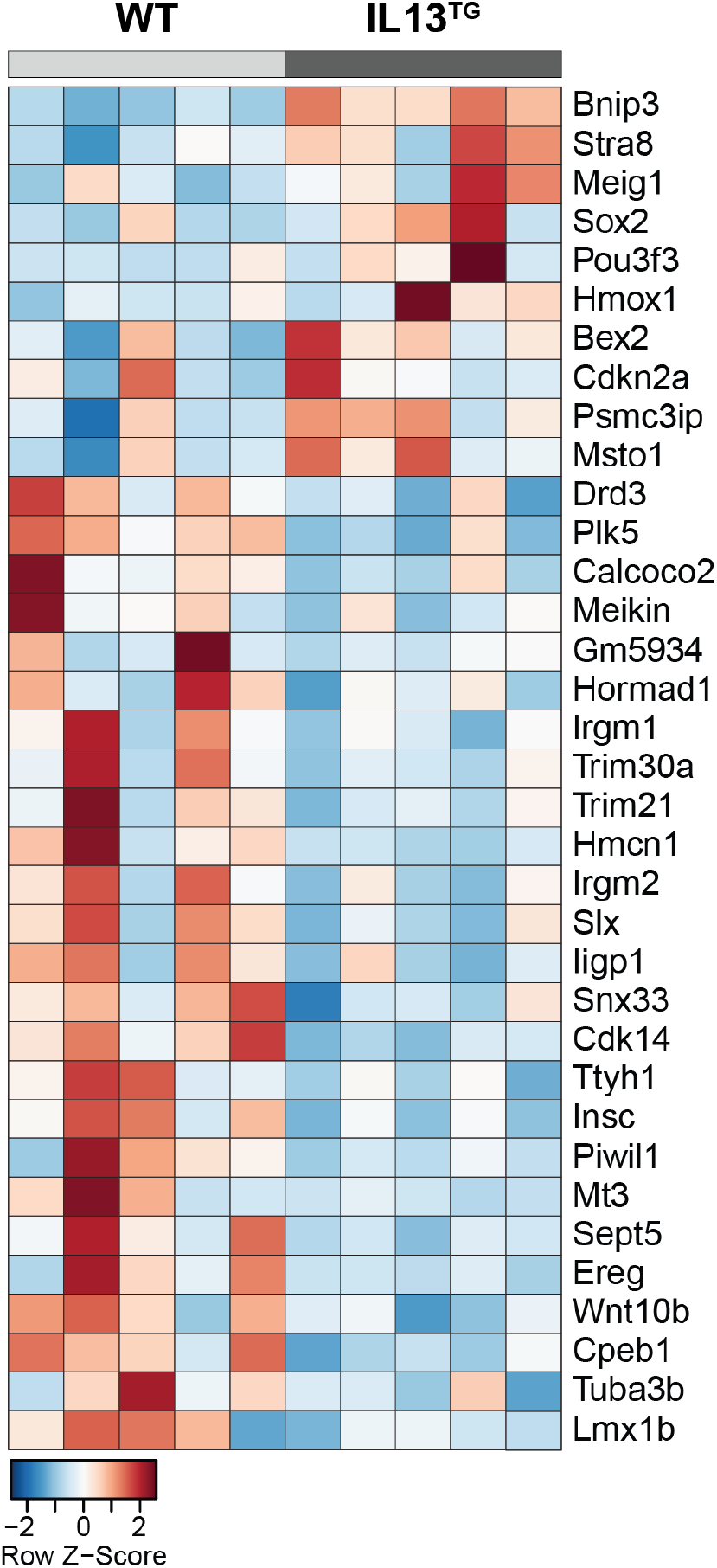
Post injury isolated muscle satellite cells’ RNA sequencing shows dysregulation of multiple transcripts related to cell cycle and autophagy in the *IL13^TG^* animal as determined by gene ontology (GO) enrichment. Heat map of gene expression shows direction of differential gene expression in *IL13^TG^* versus *WT* cells (n=5). Red indicates up-regulated gene expression and blue indicated down-regulated gene expression.

### Satellite cells from COPD mice demonstrate evidence of dysregulated autophagy

To directly investigate the potential role of autophagy dysregulation in COPD mice satellite cells, we crossed *IL13*^*TG*^ and *IL13*^*WT*^ mice with an animal expressing an enhanced green fluorescent protein (EGFP) sequence fused with the LC3 family proteins, which retain constitutive fluorescence of autophagosomes after their formation until these fuse with lysosomes [57]. In this model, autophagosome mass can be unbiasedly scored in isolated cells by determining the mean fluorescence intensity (MFI) generated by GFP puncta [32, 58]. Satellite cells isolated from COPD and wild type mice were then analyzed, which revealed an increased MFI in satellite cells obtained from COPD mice (**Figure 5**). These data strongly suggests dysregulation of autophagy in satellite cells from COPD mice, which is characterized by an increased number of autophagosomes. Importantly, this feature, which could be driven by a deacceleration of autophagy flux or increased autophagosome formation [58, 59], has been previously found in locomotor muscle biopsies from COPD patients [37, 39].

**Figure 5.**
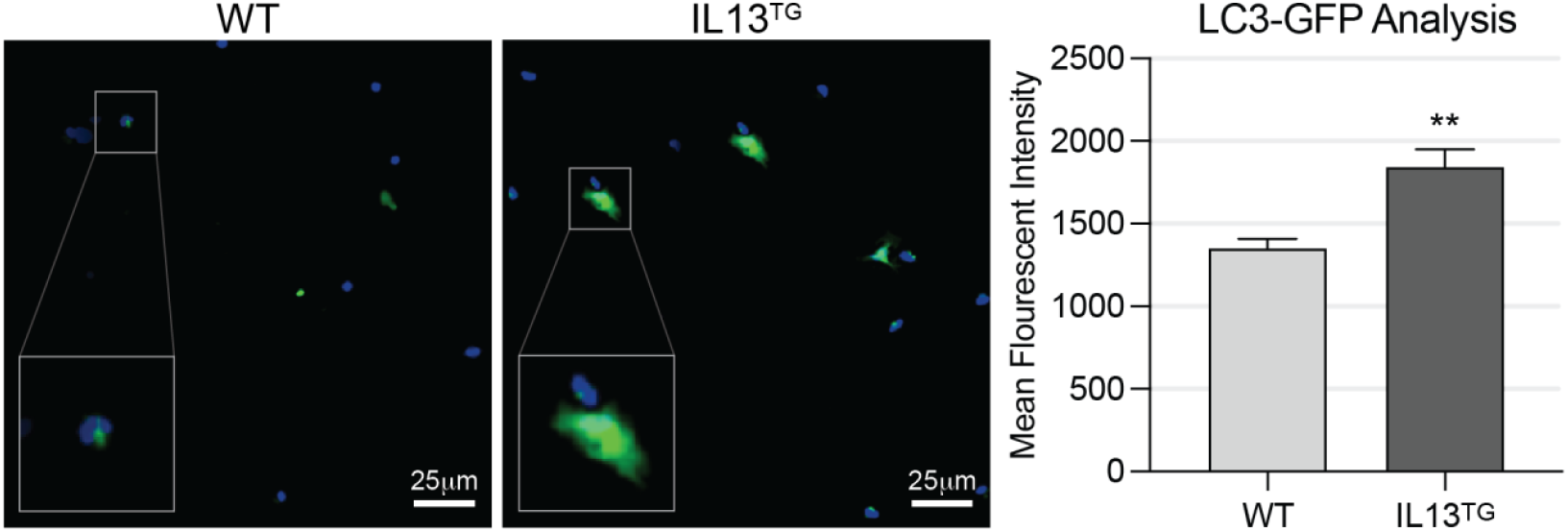
Isolated *IL13^TG^* myogenic cells exhibit dysfunctional autophagosome turnover. *LC3-GFP/IL13^TG^* reporter mice show abnormal build-up of puncta compared to *WT* mice. Mean fluorescent intensity (MFI) generated by *LC3-GFP* puncta measured in 1000 cells per animal show significantly increase MFI in *IL13^TG^* compared with *WT* counterparts, indicating a dysfunctional autophagosome turnover (n=4). Notice that these experiments were conducted without bafilomycin. \*\**P <* 0.01.

### Autophagy stimulation leads to an improved satellite cells replication capacity and post-transplant myogenesis

We then sought to determine whether pharmacologic autophagy induction can attenuate the satellite cells replicative and myogenic deficits in COPD. COPD mice are chronically debilitated [43]; thus, in order to reduce the potential bias entailed by the administration of a toxic drug due to off-target effects, we chose the lower toxicity drug spermidine, which is an inhibitor of the autophagy antagonist acetyl transferase EP300 [60] and has been previously shown to improve myogenesis [20]. To interrogate the effect of spermidine on COPD mice autophagy, we provided the drug in drinking water to LC3-GFP-expressing COPD and wild type mice. Satellite cells were later isolated and mean fluorescence intensity generated by GFP puncta was scored, which demonstrated values not significantly different among COPD and wild type mice, suggesting a normalization of autophagosome turnover (**Figure 6-A**). To define whether that effect correlated with a better replication capacity, we conducted an EdU incorporation assay. Spermidine administration led to an improvement of replication capacity of freshly isolated satellite cells from COPD mice, reflected by EdU incorporation which became not significantly different from wild type animals’ (**Figure 6-B**). To determine if spermidine leads to a better myogenic potential of satellite cells, we provided it to COPD and wild type, RFP-expressing mice and then conducted transplantation experiments using freshly isolated satellite cells as previously described. Cells obtained from spermidine-provided COPD mice recovered myogenic potential as demonstrated by red fibers development in the receiving, wild type mice (**Figure 6-C;** see cartoon above). These data suggest that the autophagy-boosting drug spermidine improves satellite cells autophagosome turnover, replication capacity and myogenic potential in an animal model of COPD. A summary of the described findings is presented in **Figure 7**.

**Figure 6.**
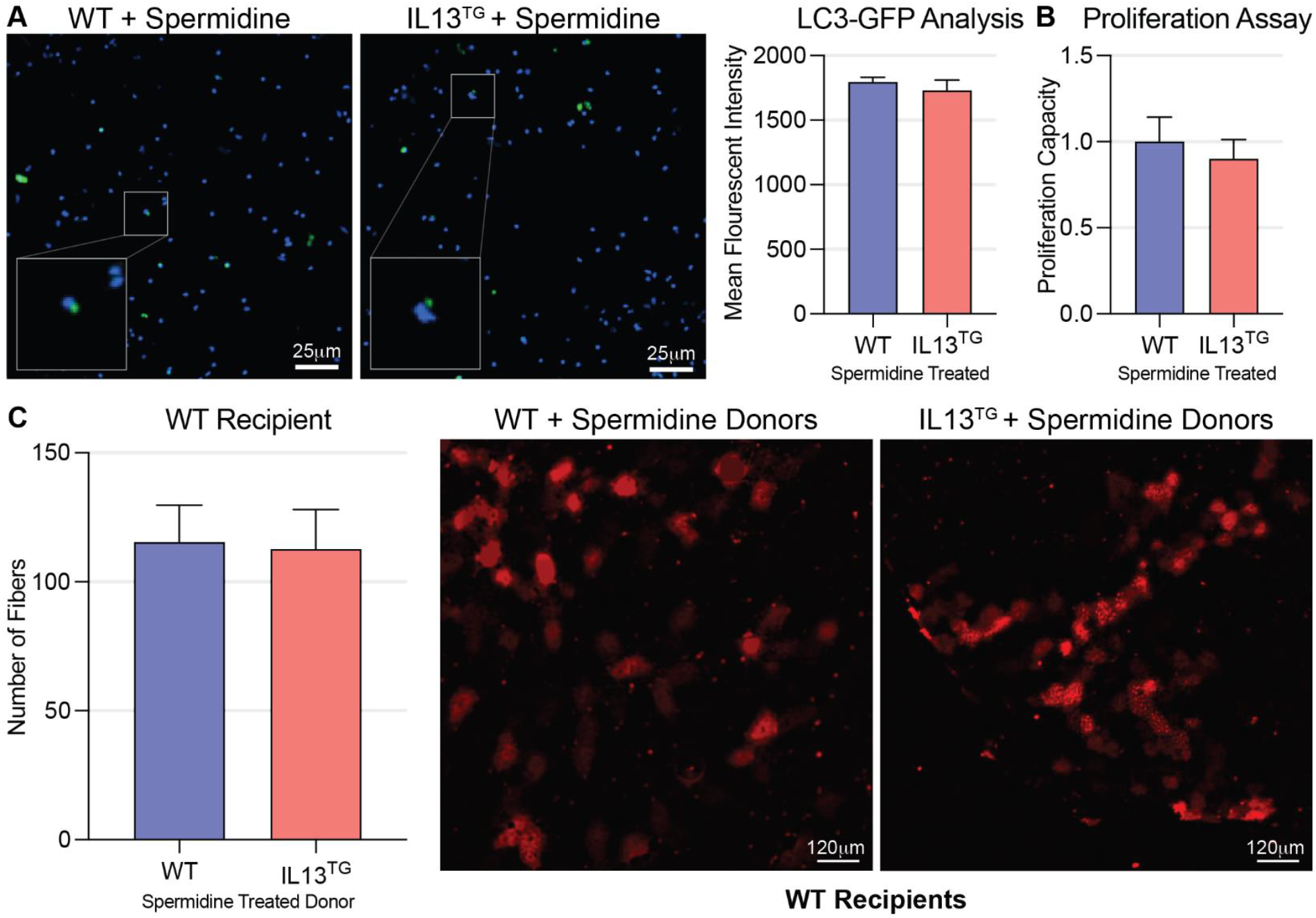
Spermidine-induced autophagy improves myogenesis in COPD mice. **A:** *LC3-GFP/IL13^TG^* reporter mice show normalization of puncta buildup and MFI in isolated satellite cells after treatment with spermidine (n=4). **B:** EdU proliferation assay shows no difference between *IL13^TG^* and *WT* animals after treatment with spermidine (n=4). **C:** Transplantation experiments in *IL13^WT^* recipients conducted as previously described show no difference in number of recovered red fibers when both *IL13^TG^* and *IL13^WT^* donors were treated with spermidine (n=4).

**Figure 7.**
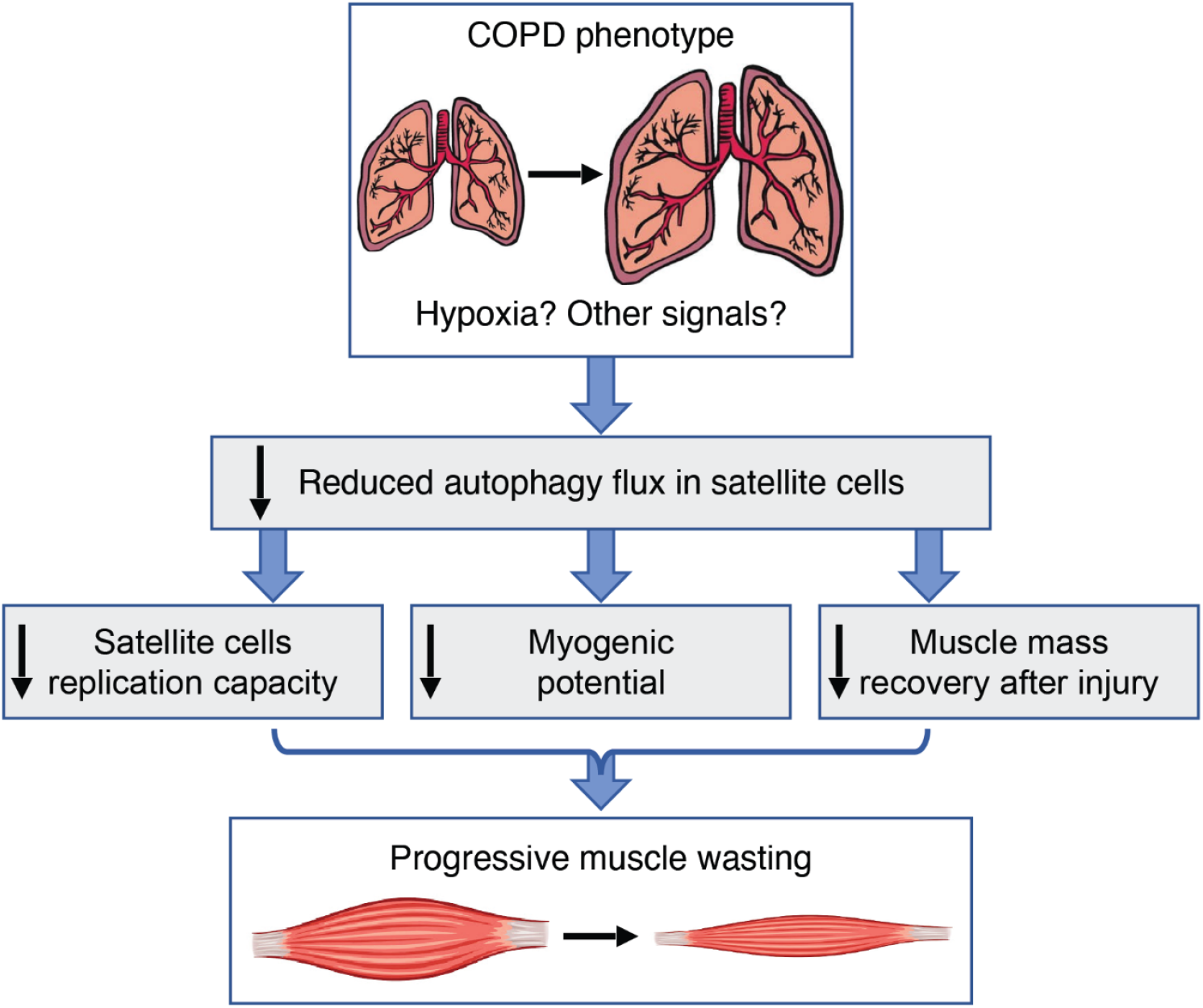
A proposed schematic model showing that the COPD lung phenotype leads to altered autophagy flux which triggers myogenic dysfunction and undermines muscle repair, contributing to muscle wasting over time

## Discussion

While autophagy dysregulation is conspicuous in skeletal muscle biopsies from COPD patients [37, 39], there is debate on the implications of that abnormality on muscle dysfunction [38]. Indeed, while autophagy activation is needed for muscle integrity preservation [61], it has also been associated with muscle loss [40]. For instance, it remains unclear whether the abnormal autophagy signature from COPD patients muscles reflects increased or reduced autophagosome flux [38]; and, more fundamentally, if it plays a role in the pathophysiology of the COPD myopathy or instead it represents an inconsequential hallmark of that process. Myogenesis, which contributes to skeletal muscle integrity [49], depends in part on autophagy regulation, which provides the bioenergetic needs for myogenic satellite cells activation and prevents their transition into senescence [17, 20, 32-36]. We report here that myogenic capacity and satellite cells replication are both reduced in an animal model of COPD. Although relatively unexplored in non-primarily muscle diseases, myogenesis is emerging as an important mechanism regulating muscle integrity in COPD and other pulmonary diseases [18, 21-27]. Following an injurious event, satellite cells undergo an initial phase of symmetrical replication giving rise to similar daughter cells which expand the pool of myogenic cells [62]. After that, asymmetric division in an apical-basal orientation gives rise to two daughter cells, only one of which activates the transcription factor Myf5 and commits to myogenic differentiation [8]. Myogenesis is highly influenced by the interaction between progenitor cells and their environment including the muscle stem cell niche [8]. Our combination of satellite cell transplantation and lineage tracing assays supported a dominant role of intrinsic stem cell dysfunction, which led to RNAseq analysis of these cells that revealed dysregulation of multiple candidate genes potentially regulating the observed replicative dysfunction. Indeed, we found various genes related to autophagy and cell division to be dysregulated in the COPD animal satellite cells, which were investigated with specific surrogates. LC3 family proteins involving MAP1-LC3, GATE-16, and GABARAP, which are mammalian counterparts of yeast Atg8, are present on complete autophagosomes and degraded after autophagosome fusion with lysosomes [63]. We used a green fluorescent protein (GFP)-fused LC3-reporting animal to track autophagy turnover on freshly isolated satellite cells, which demonstrated an increased mean fluorescence intensity in cells procured from the COPD animal in comparison with wild type counterparts [58]. That observation could represent either an acceleration of baseline autophagosome generation or reduced autophagosome-lysosome fusion leading to an accumulation of GFP-LC3 signal. Remarkably, a similar signature of accumulated autophagosomes has been described in muscle biopsies from COPD patients [37, 39]. Moreover, in COPD patients, the amount of accumulated autophagosomes was found to be correlated to the severity of lung dysfunction as reflected by the reduction in the forced expiratory volume in the first second (FEV1) [37].

To further determine whether autophagy dysregulation in satellite cells contributed to a reduced proliferative capacity and myogenic potential, we treated mice with spermidine, a drug known to upregulate autophagy flux and myogenesis [20, 36]. Although spermidine induces autophagy via the inhibition of acetyltransferase EP300 [60, 64], and does not affect the phosphorylation of mTOR or its substrate ribosomal protein S6 kinase [65], the specific mechanism of spermidine-induced autophagy in COPD mice will be a matter future research. Given that autophagy is also dysregulated in the muscle fiber, the beneficial effect of spermidine could, at least partially, occur indirectly via its effect on autophagosome turnover at the non-satellite cell level. Future studies using a satellite cells specific autophagy knockout animal [20] will address whether autophagy integrity in the satellite cell compartment is required for spermidine or other drugs to cause improved myogenesis in pulmonary emphysema. Importantly, spermidine is remarkably less toxic and immunosuppressive than rapamycin [66], making it an attractive target to investigate its potential myogenic benefits in chronically debilitated COPD patients.

While most of the research conducted in the field of COPD-associated muscle dysfunction has so far focused on the conspicuous reduction of muscle mass [1,4-6, 43, 44, 47, 67-69], to our knowledge this is the first mechanistic study that disaggregates the investigation of autophagy-driven myogenesis in the context of pulmonary emphysema. The fact that the MMP1 model of COPD, which is bred on a non-C57 background and develops the COPD phenotype via different mechanisms [51], also demonstrates skeletal muscle dysfunction and reduced satellite cells replication suggests that it is the pulmonary emphysema *per se* that drives the myogenic dysfunction. It is important to emphasize that the MMP1 holds a constitutive transgene, and thus myogenesis and muscle development could be disrupted before the COPD phenotype fully develops [46]. Due to this important limitation, the MMP1 line serves only as a validation, but not the primary focus, of the explored mechanisms.

The upstream mechanism regulating dysfunctional autophagy in this model remains to be elucidated. While *IL13*^*TG*^ mice develop conspicuous hypoxemia, it has recently reported that exposure of myoblasts to hypoxia leads to an increase in histone deacetylase C9 (HDAC9) expression, which deacetylates H3K9 of autophagy-related genes and thus epigenetically inhibits autophagosome formation [70]. Low oxygen could also contribute to dysfunctional myogenesis via hypoxia inducible factor 1α (HIF1α), which inhibits canonical Wnt signaling and undermines satellite cell function that way [71]. Future studies can address these and other hypotheses. Interestingly, while downregulation of muscle succinate dehydrogenase (SDH) has been consistently shown in COPD patients [72-74], we have recently reported that reduced expression of SDH subunit C (SDH-C) contributes to muscle metabolic dysfunction in *IL13*^*TG*^ mice [48]. SDH dysfunction leads to succinate accumulation [48], which has recently been shown to influence the regulation of cyclin-dependent kinase 2 (CDK2)-driven phosphorylation in satellite cells via succinate receptor SUCNR1 [75]. As CDK2 is a critical regulator of cell division, succinate could articulate metabolic and myogenic dysregulations in this context.

Importantly, the observed COPD-induced dysfunctional myogenesis demonstrates distinct features in comparison to other non-primarily muscular diseases. For instance, cancer cachexia has been recently shown to cause dysfunctional myogenesis via NF-kappa B-mediated Pax7 dysregulation, which is associated with an increased replication of satellite cells and altered differentiation [18], in contrast to our findings of reduced satellite cells replication. Also, while the number of myogenic cells in the COPD animal is not significantly different from the wild type counterparts, dysfunctional myogenesis associated with advanced age is associated with a p16INK4a-regulated transition from quiescence to senescence with a reduction of satellite cells number [17]. A recent report also suggests a reduced number of satellite cells in skeletal muscles from patients surviving critical illness and developing muscle wasting [21, 76].

The mechanism articulating autophagy dysregulation with reduced satellite cell replication and myogenesis in pulmonary emphysema remains to be elucidated. Previous data indicates that inhibition of autophagy suppresses ATP levels needed for satellite cells activation, which is partially rescued by exogenous pyruvate supply as an energy source [32]. The interaction between mitochondrial integrity and autophagy flux has been demonstrated in other models [34-36] and could also be relevant here as well. Future studies analyzing the metabolic substrate alterations in COPD mouse satellite cells could provide relevant data in that regard.

We realize this study has the limitation of being based on a murine lung-specific gene over expression leading to pulmonary emphysema phenotype. It is important to emphasize that while IL13 regulates the pathophysiology of pulmonary emphysema [77] and correlates with disease severity in that setting [78], IL13 is not described to cause muscle toxicity independently of the COPD phenotype. Moreover, while IL4/IL13 signaling has been found to promote muscle trophism and not dysfunction [79, 80] at the expense of higher myogenic capacity, this animal model demonstrates reduced satellite cell replication rate, suggesting that IL13 is unlikely to be associated with the reported myogenic phenotype. Indeed, this model does not demonstrate significant muscle loss until the pulmonary emphysema develops, at 8 weeks post transgene induction start [43, 44]. However, the possibility muscle toxicity driven by IL13 independently of the pulmonary emphysema development cannot be fully excluded, as it has recently been reported on lungs stem cells activity [81]. **Conclusion:** Skeletal myogenesis is dysfunctional in animal model of COPD, which is associated with dysregulated autophagy in satellite cells and attenuated by systemic administration of spermidine.

## Methods

### Animals

#### COPD model (Primary)

Experiments were conducted using CC10-rtTA-IL-13 (*IL13^TG^*) doxycycline-inducible transgenic mice that develop chronic lung remodeling reminiscent of pulmonary emphysema upon induction. CC10-rtTA-IL-13 heterozygote animals were bred to C57BL/6 wild-type (WT) mice to obtain hemizygous *IL13*^*TG*^ experimental animals and homozygous *IL13*^*WT*^ littermate controls. Both *IL13*^*TG*^ (emphysema) and *IL13*^*WT*^ (non-emphysema, used as control littermates) mice were provided doxycycline in the drinking water along with sucrose, starting at 5 weeks of age for a total of 17 weeks (∼22 weeks of age). Both male and female mice were used for the studies and results are reported as aggregate data from both sexes. Because female mice have been shown to have higher muscle regenerative capacity [82], to prevent a sex-specific biasing we normalized the COPD animal EdU incorporation assay as the percentage of the wild type same sex counterpart, and conducted the transplantation assays using same sex donors and using, in the same receiving animal, one TA muscle for COPD-and the contralateral TA for wild type-donated cells. Food and water were accessible ad libitum and a 12-hour light/dark cycle was maintained by our Animal Resource Facility (ARF). Samples were collected directly after euthanasia by cervical dislocation. The time elapsed between animals’ euthanasia and sample procurement never exceeded 3 min when methodologically feasible.

#### COPD model (Validation)

Six lung cell-derived MMP1 transgenic mice (22 weeks old), generated as previously described [51], were used directly upon import from Columbia University for specific characterization of muscle function readouts and EdU incorporation assay.

#### βActin-DsRed

Transgenic mice expressing red fluorescent protein (hence named “*Red^+/+^*”) variant DsRed.MST under the control of the chicken β-actin promoter coupled with the cytomegalovirus (CMV) immediate-early enhancer [55] were purchased from the Jackson Laboratory (Stock No: 005441) and used for transplantation experiments in the hemizygous form (hence named “*Red^+/-^*”).

#### GFP-LC3

Mice expressing GFP (EGFP)–LC3 cassette inserted between the CAG promoter (cytomegalovirus immediate-early (CMVie) enhancer and chicken β-actin promoter) on a C57BL background were purchased from RIKEN Bio-Resource Center, Japan (#BRC00806). Genotype was determined as previously reported and confirmed by direct visualization of the GFP puncta [58].

#### GFP-Pax7

An animal expressing a IRES-CreERT2 fusion protein downstream the Pax7 stop codon (Jackson Lab, Stock No: 017763) was crossed with a loxP-flanked STOP cassette holding animal (Jackson Lab, Stock No: 007906) to obtain a double homozygous animal (*Pax7-Cre^+/+^Lox-GFP^+/+^*). The double homozygous animal was then crossed with the hemizygous *IL13^TG^* animal to obtain both *IL13^TG^* and *IL13^WT^ GFP-Pax7* reporting littermates for experiments. This tracing method has been shown to capture 95% Pax7-expressing cells in skeletal muscle [53].

### Oxygen Saturation Measurement

At 17 weeks post-doxycycline initiation (22-23 weeks of age), oxygen saturation of *IL13^TG^* and *WT* animals was measured using the MouseOx Plus pulse oximeter (Starr Life Sciences Corp., Oakmont, PA) with the small sized collar sensor as recommended by the manufacturer. In short, hair was carefully removed from the animal’s neck using hair remover (Nair^®^) and the collar was used to take an unanesthetized saturation measurement.

### Lungs procurement and histology

Animals were sacrificed via cervical dislocation; a median sternotomy was performed immediately to collect lungs. Lungs were fixed overnight in 10% formalin, embedded in paraffin, cut at 5 μm-thick sections, and stained with hematoxylin and eosin (H&E). Sections were imaged using a Cytation 5 imager (BioTek, Winooski, VT)

### Muscle procurement and determination of muscle mass

At 17 weeks post doxycycline initiation (22-23 weeks of age), immediately after animal euthanasia, leg skin was dissected with tweezers and muscles exposed. Under real time magnification using an illuminated magnifier (Omano, China), muscles were dissected individually, cutting first the distal tendon and gently removing the entire muscle using a dissecting scissors (Roboz RS-5840). After the proximal tendon was severed, the muscle was put in contact with a gauze under magnification to eliminate remaining blood, fat, and if needed also remaining tendon using a scissor. Muscle was then weighed using an analytical balance (Sartorius Entris, Germany). Mass determinations were done on gastrocnemius and tibialis anterior (TA) muscles.

### Isolated Muscle Contractility

Extensor digitorum longus (EDL) muscles were surgically isolated from the mouse by carefully tying a suture around the tendon at each end and cutting the tendons to release the muscle. Special care was taken not to stretch or damage the muscle integrity as previously established [83]. Once removed, the analyzed muscle was equilibrated for 15 minutes in ice cold Ringer’s solution supplemented with 5.5 mM glucose, adjusted to a pH between 7.4 and 8.0, and slowly bubbled with carbogen. The muscle was then suspended between the isometric force transducer (Harvard Apparatus, Holliston, MA) and the platinum stimulating electrode tissue support (Radnoti, Covina, CA, 160152), lowered into the 25 mL tissue bath (Radnoti, Covina, CA, 166026) containing the same solution, also bubbled slowly with carbogen but this time at room temperature; and allowed to equilibrate for an additional 15 minutes. Muscle tension was escalated until baseline tension started to increase. A single 1 Hz 40-volt stimulus was delivered with a Grass S-88 electrical stimulator and the peak contraction was recorded. After a 30 second rest, voltage was increased by 10v and delivered again, recording peak contraction. This process was repeated until no additional increase in the peak contraction force was observed. The optimal length of the muscle was then determined by slowly increasing the muscle tension and delivering a single, maximal stimulus as previously determined above, while recording the peak force. After a 30-second rest, tension was slightly increased, and another stimulus was delivered while recording peak contraction. This process was repeated until maximal peak contraction force was achieved. Subsequent stimuli were delivered at 1, 10, 20, 30, 50, 80, 100, and 120 Hz while recording the peak force at each point and allowed for 1-minute rest between each stimulus.

### Grip Strength Determination

Animal’s four limb grip strength was determined using a GSM Grip Strength Meter (Ugo Basile, Gemonio, Italy, 47200) with the full grasping grid. The mouse was help by the tail and placed on the grasping grid, once the test was started pressure was steadily applied to the mouse’s tail until failure and release from the grasping pad. The peak force was recorded, and the mouse was given a 1-minute rest, this was repeated for a total of 5 replicates.

### Muscle Fiber Cross Sectional Area Measurement

Muscle fiber delineations at 10X magnification on sections stained for Laminin were analyzed with CellProfiler software (Broad Institute, Cambridge MA) to measure fiber cross sectional area in an unbiased manner. Output muscle traces were reviewed to assure accuracy of measurement of the software pipeline.

### Muscle Immunofluorescence

Muscle sections were fixed for 15 minutes in acetone at -20°C, and then left at room temperature to dry for 30 minutes. Blocking was performed using Mouse-on-mouse blocking reagent (Vector Labs, Burlingame, California) for 1 hour at room temperature. Sections were then incubated for 45 minutes at 37°C with the primary antibody at 1:100 for anti-laminin (L9393, Sigma-Aldrich). Three washes were then performed with PBS. Secondary antibody for anti-rabbit IgG-Alexa 594 (Jackson ImmunoResearch Laboratories) was added at 1:250 for 45 minutes at 37°C. Three washes were then performed with PBS. Samples were mounted with Ibidi Mounting Medium (Martinsried, Germany). Images were captured the same day using confocal microscopy (Leica SPE).

### RNA Extraction, cDNA Synthesis, and Quantitative RT-PCR

RNA from tibialis anterior (TA) muscle was extracted using NucleoSpin RNA kit (Machery-Nagel, Düren, Germany). cDNA was synthesized using Quantitect Reverse Transcriptase Kit (Qiagen). Quantitative RT-PCR was performed using iTaq Universal SYBR Green Supermix (Bio-Rad) on a CFX96 Real-time PCR detection system (Bio-Rad). Each sample was run in triplicates, and relative expression levels of transcripts of interest were calculated using the comparative Ct (ΔΔCt) method with glyceraldehyde-3-phosphate dehydrogenase (GAPDH) as a housekeeping gene. Primers were purchased from Integrated DNA Technologies (IDT, IA), and a list of their sequences is presented in **Supplementary Table 3**.

### Muscle Histology

At 23 weeks of age, freshly isolated muscles were placed on saline-moistened gauze in a 60-mm culture dish on ice until freezing. A metal cup containing isopentane was cooled in liquid nitrogen until crystals formed of isopentane at the bottom of the cup. Muscles were transferred to precooled Tissue-Tek embedding cassettes (EMS, Hatfield, PA; 62520), which were dropped into the cooled isopentane, submerging the muscle for 1 min. Muscle samples were then drained and dried on gauze pads at -20°C to remove all isopentane. Frozen muscles were adhered to the sample stage using a small amount of Tissue-Tek optimal cutting temperature (OCT) compound (EMS, Hatfield, PA, 62550) and 10-micron sections were taken for analysis using a Leica CM1860 Cryostat (Wetzlar, Germany).

### Muscle Fiber Cross-Sectional Area

Measurement Muscle fiber delineations at 10X magnification on sections stained for laminin were analyzed with CellProfiler software (Broad Institute, Cambridge, MA) to measure fiber cross sectional area in an unbiased manner. Output muscle traces were reviewed to assure accuracy of measurement of the software pipeline.

### Induction of Muscle Injury

Mice were anesthetized with isoflurane and hair was removed from the hind legs using Nair^®^. The right tibialis anterior (TA) muscle of the hind leg was injected with 50uL of sterile saline while the left TA muscle was injected with 50uL of 1.2% barium chloride in sterile saline as previously established [56]. Mice were left to recover in their cage while closely monitored. Twenty-one days after the initial injury, a second injury was performed using the same protocol. At specified days after the second injury procedure, the TA muscles were collected and frozen histologically. Due to the COPD-induced muscle atrophy that is intrinsic to the animal model [43, 44, 47, 48], we scored the fibers cross sectional area after full reconstitution of histological integrity, in the left TA muscle, as a percentage of the expected value, which is the CSA of the right (saline-injected) TA muscle.

### Satellite Cell Isolation

Skeletal muscles were collected from the hind legs of the mice and cellular dissociation was performed. Mouse skeletal muscle satellite cells were first enriched using a Mouse Satellite Cell Isolation Kit (MACS Miltenyi Biotech, 130104268) and then selected for using anti-integrin α-7 microbeads (MACS Miltenyi Biotech, 130104261) to insure population purity as previously established [52].

### FACS Isolation and RNA Sequencing

Cardiotoxin was chosen for this experiment rather than BaCl_2_ as a more complete and robust injury was required in multiple leg muscles, and a larger volume of drug was required. Indeed, when the volume of BaCl_2_ needed was used for this experiment, the animals did not survive presumably due to the pre-existing lung condition and the known respiratory muscle paralysis caused by barium [84]. Muscle injury was caused on fully induced *IL13^WT^* and *IL13^TG^* mice three days prior to cell collection by injecting 10uM cardiotoxin (Millipore Sigma, 217503). Injections were performed in 10uL aliquots along the muscle body -each tibialis anterior (TA) muscle was injected with 40uL total and each gastrocnemius (GN) muscle was injected with 80uL total. Three days after injury all hind leg muscle was collected and enzymatically dissociated and filtered. Filtered cells were resuspended in 300uL of buffer (3mM EDTA, 5% FBS) and the following antibodies were added; 10uL ITGA-7 Alex 647 (AbLab Custom, Canada), 15uL CD34 Biotin (Miltenyi, Germany), 2uL Sca-1 (eBiosciences, USA), 2uL CD31 (eBiosciences, USA), 2uL CD45 (eBiosciences, USA), 2uL CD11b (eBiosciences, USA). After 30 minutes on a rocker at 4°C, cells were washed and centrifuged at 250 RCF for 5 minutes at 4°C, supernatant was removed, cells were resuspended in 250uL of buffer, 4uL of PE-Cy7 streptavidin secondary antibody (Invitrogen, USA) was added and incubated for 20 minutes on a rocker at 4°C. Cells were then washed with buffer one additional time and filtered for sorting with our BD FACSAria™ II. Collected cells were centrifuged and buffer was removed as previously described. RNA was isolated using the Arcturus PicoPure Isolation Kit (Applied Biosystems, KIT0204) following manufacturer’s protocols. Low-input library preparation was performed using the SMART-Seq^®^ Stranded Kit (Takara, 634443) following manufacturer’s suggested protocols. Sequencing was performed on an Illumina HiSeq 4000 instrument at The Stanford Center for Genomics and Personalized Medicine, Stanford University; and differential analysis was performed at The University of Pennsylvania Bioinformatics Core.

### EdU Proliferation Assay

Cells, isolated as previously described, were plated on 20mm insert glass bottom plates (MatTek, P35G1.520C) with growth media containing 0.05 mM EDU. The media was changed carefully every 16 hours until the cells were fixed at 40 hours using 4% paraformaldehyde for 30 minutes at 4°C. EdU incorporation was visualized with a Click-iT kit (Thermo-Fisher Scientific, C10337) following the manufacturer’s instructions and as previously established. Spermidine experiments were designed similarly however, animals were provided 3mM spermidine for 4 weeks in drinking water until collection.

### Satellite cells Transplantation

One day before the transplantation, injury was performed on recipient animals by removing the hair from the hind legs and injecting each of the animal’s TA muscle with 25uL of 1.2% barium chloride. Simultaneously, animals began receiving 15mg/kg Cyclosporine A by subcutaneous injection (Novartis, East Hanover, NJ) which was continued every day until the muscle collection date and provided to increase the allotransplanted cells viability. Integrin α-7 enriched satellite cells were collected from donor mice as previously described, and 50,000 cells were immediately injected into each TA of the recipient mouse: one TA received cells from an experimental donor while the contralateral leg, used as an internal control, received cells from a control animal. Two weeks after the transplant, TA muscles were collected and histologically frozen. Fresh frozen sections were obtained, and images were immediately acquired using a Leica DM-IRB fluorescent microscope; and total numbers of auto-fluorescent red cells were counted. Spermidine experiments were designed similarly however, donor animals were provided 3mM spermidine for 30 days in drinking water until collection.

### Pax7-GFP Cell Counting

Mice were bred with an inducible Pax7-GFP reporter (Pax7-Cre^+/-^Lox-GFP^+/+^*IL13^TG^*) and collected at 17-weeks post-doxycycline initiation (∼22 weeks of age). Age and gender matched control mice (Pax7-Cre^+/-^Lox-GFP^+/+^*IL13^WT^*) were also collected at 17-weeks post-doxycycline initiation (∼22 weeks of age). Mice were induced with 500 mg/kg tamoxifen chow (Envigo, TD.130857) for two non-consecutive weeks during the final 30-day period of doxycycline. Mice were euthanized and TA muscles were procured histologically as previously described. GFP-positive Pax7 cells were counted per section automatically using a Cytation5 cell imager (Biotek, Winooski VT).

### LC3-GFP Autophagy Assay

Mice were bred with a constitutively active LC3-GFP reporter (LC3-GFP^+/+^*IL13^TG^*) and collected at 17-weeks post-doxycycline initiation (∼22 weeks of age). Age and gender matched control mice (LC3-GFP^+/+^*IL13^WT^*) were also collected at 17-weeks post-doxycycline initiation (∼22 weeks of age). Satellite cells were isolated from the hind legs of these mice as previously described. Mean fluorescence intensity (MFI) was measured automatically using a Cytation5 cell imager (Biotek, Winooski VT). Due to the heterogeneity produced by this method on individual cells, we measured the MFI of about 1000 isolated cells per animal to generate an average LC3-GFP expression. A primary mask was set using the nuclei and a secondary mask was constructed by adding a known perimeter to the nuclear mask to include the cytosol. MFI readings were taken as an aggregate of both masks. The images chosen for publication are representative of these average data. Spermidine experiments were designed similarly however, animals were provided 3mM spermidine for 30 days in drinking water until collection.

### Drug Preparations

Both *IL13^WT^* and *IL13^TG^* animals were given water with 0.5 mg/mL doxycycline and 0.5 g/mL sucrose starting at age 5 week for a total of 17 weeks. Spermidine was delivered to animals at 3mM in the drinking water in addition to the doxycycline and sucrose when required for a total of 4 week. Tamoxifen was delivered to animals when required in the chow at 80 mg/kg (Envigo, TD.130857), which was given to the animals for two non-consecutive weeks and stopped at least one week before any experiments.

### Statistics

Data are expressed as the means ± SE. When results were compared with a reference value, we used a single sample *t* test; when comparisons were performed between two groups, significance was evaluated by a Student’s *t* test, and when more than two groups were compared, ANOVA was used followed by the Dunnett test using GraphPad Prism software. Results were considered significant when *P*<0.05.

### Ethical approval

All the procedures involving animals were approved by the Albany Medical College Institutional Animal Care and Use Committee (IACUC protocol 18-10001), and animals were handled according to the National Institutes of Health guidelines. All methods were performed in accordance with the relevant guidelines and regulations, as stated by the Journal and public agencies.

## Data Availability

The datasets generated and analyzed during the current study will be publicly available as soon as a GEO accession number is generated (pending).

## Authors contributions

JB, LAD, DVS, and AJ designed and performed experiments; JB, JD, CEV, JAE, CGL, JD, HAS and AJ designed the experiments and wrote the current manuscript.

## Acknowledgments

Part of the results reported herein have been funded by NHLBI of the National Institutes of Health under the award number K01-HL130704 (AJ), and by the Collins Family Foundation Endowment (AJ); NIH/NHLBI R01HL086936 (JD); NIH/NHLBI 5R01HL049426 (HAS); NIH/NHLBI PO1 HL114501(JAE); R01 HL115813 (CGL).

## Supplementary material

**Figure S1.**
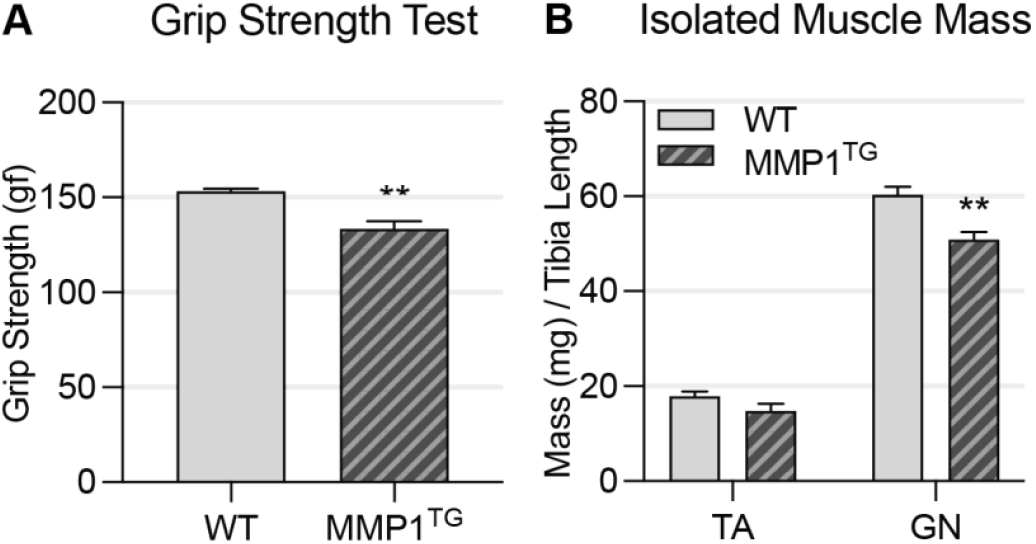
*MMP1^TG^* mice show evidence of muscle dysfunction. **A:** *MMP1^TG^* mice have significant reduction in grip strength when compared to *WT* littermates (n=4). **B:** Isolated GN muscles from *MMP1^TG^* mice have significantly decreased weight compared to *WT* mice (n=4); *\*\*P* < 0.01.

**Figure S2.**
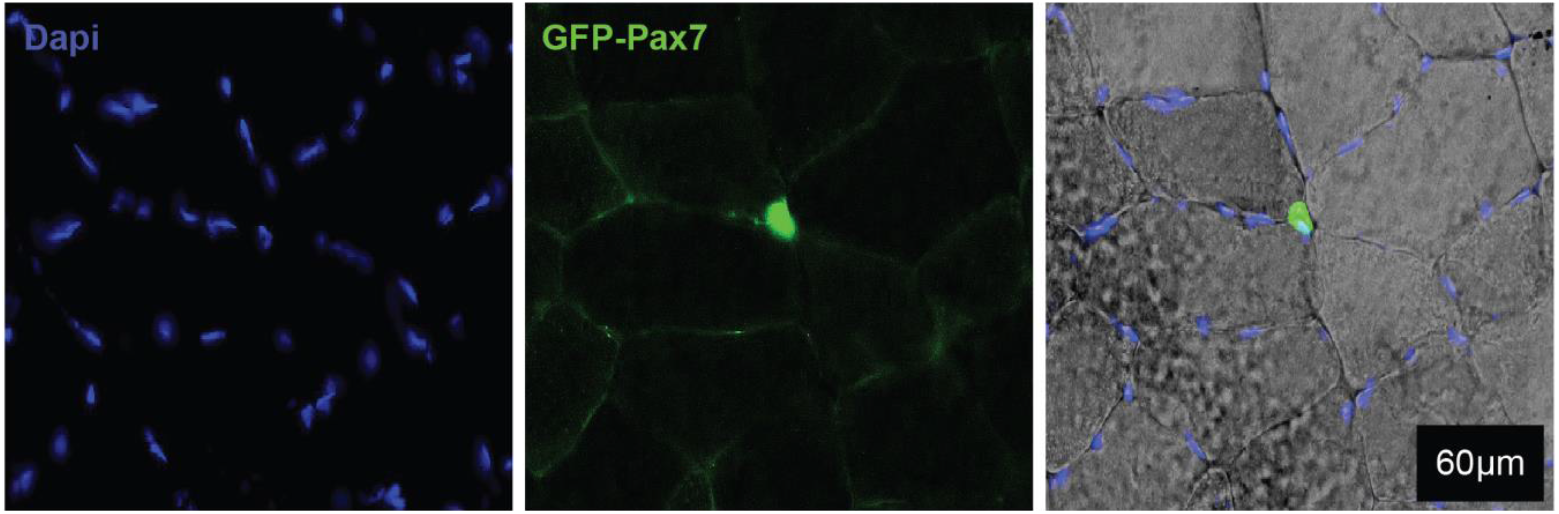
Example of GFP-Pax7/IL13^TG^ reporter mouse used for quantification of satellite cells per muscle section.

**Table S1.**
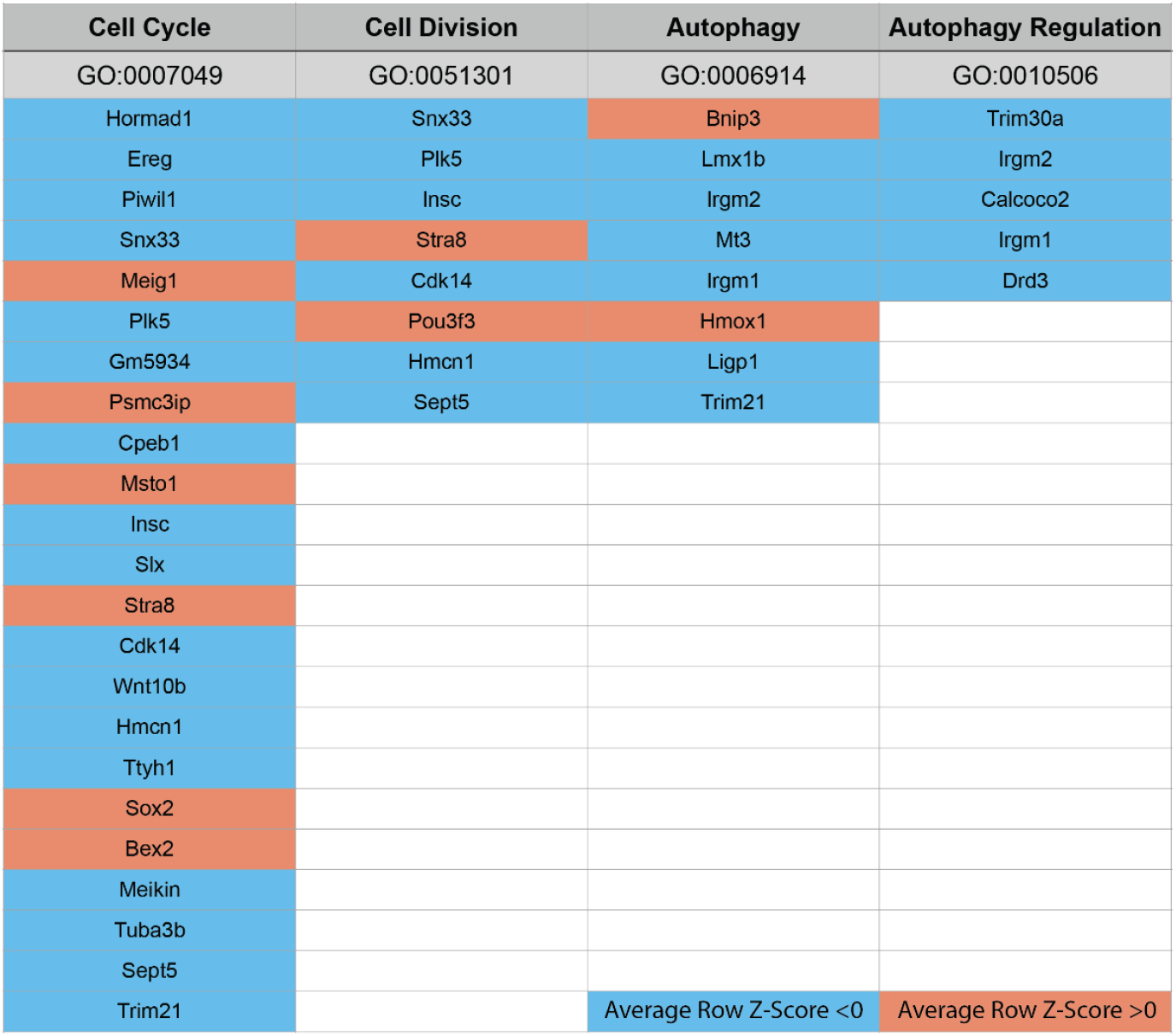
Differentially expressed genes of interest identified in RNA-Seq experiment listed under corresponding gene ontology term.

**Table S2.**
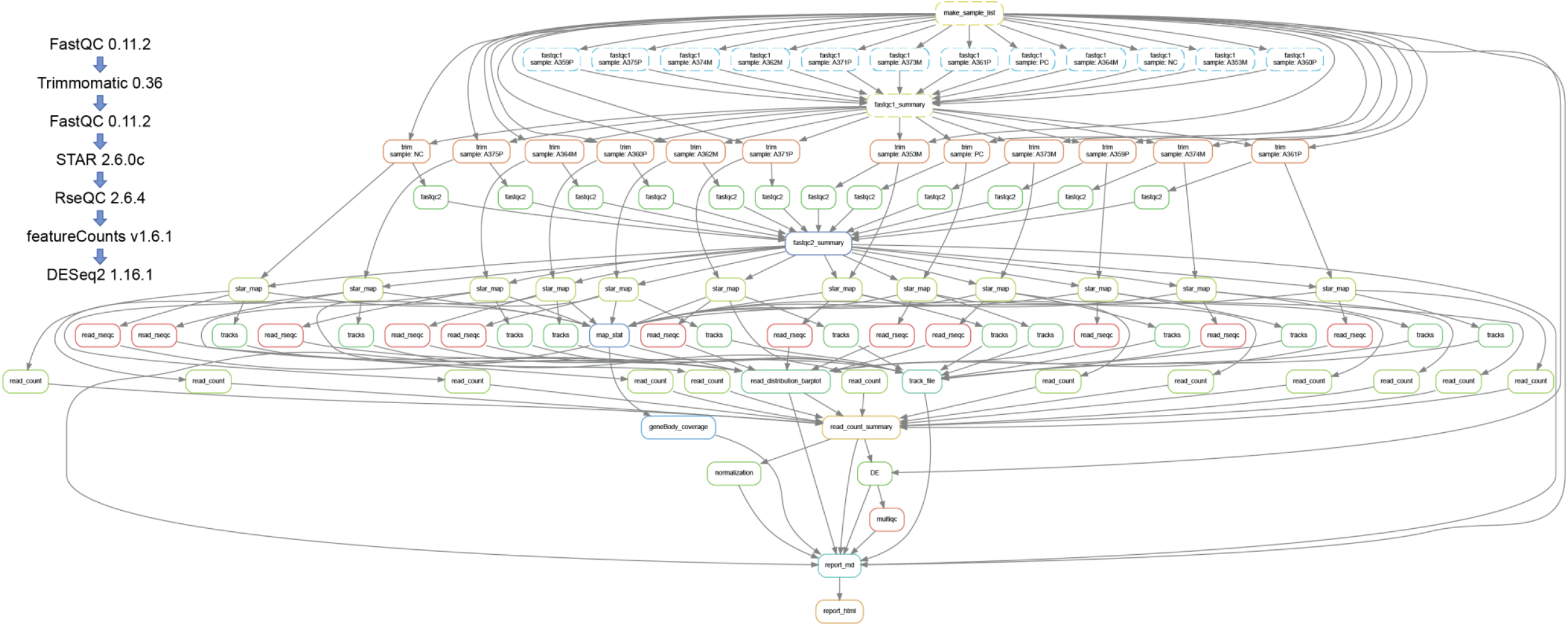
Bioinformatic pipeline used for the RNA seq analysis. Raw data will be available via a GEO accession number.

**Table S3.**
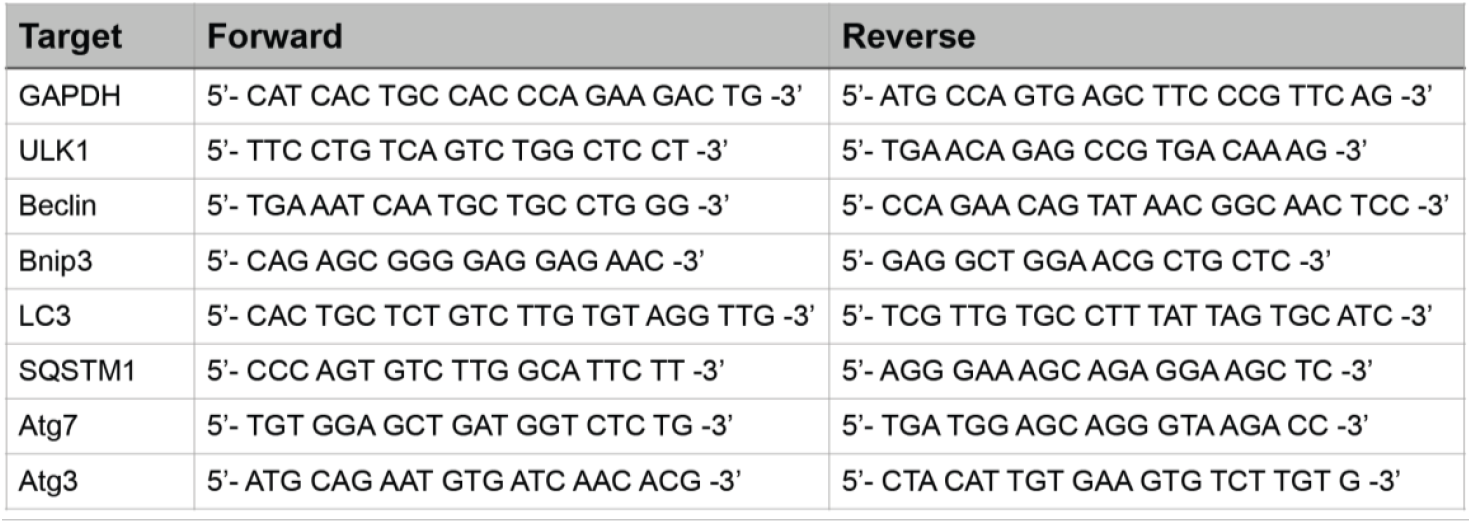
List of mouse primers used for RT-PCR

## Notes

**Conflict of Interest Statement**: The authors declare no conflicts of interest

### Competing Interest Statement

The authors have declared no competing interest.

